# Microbial signatures define the ecosystem functions of the pelagic microbiome in a basin-scale, Southwest Atlantic Ocean

**DOI:** 10.1101/2025.03.17.643744

**Authors:** Natascha M. Bergo, Francielli Vilela Peres, Danilo Candido Vieira, Flúvio Mondolon, Julio Cezar Fornazier Moreira, Rebeca Graciela Matheus Lizárraga, Amanda Goncalves Bendia, Leandro Nascimento Lemos, Alice de Moura Emilio, Augusto Miliorini Amendola, Diana Carolina Duque Castano, Mateus Gustavo Chuqui, Fabiana da Silva Paula, Renato Gamba Romano, William Soares Gattaz Brandão, Gustavo Fonseca, Daniel Moreira, Célio Roberto Jonck, Ana Tereza R Vasconcelos, Frederico P. Brandini, Vivian H. Pellizari

**Author notes:** Corresponding author Correspondence to Natascha Menezes Bergo, Francielli Vilela Peres and Vivian H. Pellizari. Natascha M. Bergo – Francielli Vilela Peres - Flúvio Mondolon - Julio Cezar Fornazier Moreira - Rebeca Graciela Matheus Lizárraga - Amanda Goncalves Bendia - Alice de Moura Emilio - Augusto Miliorini Amendola - Diana Carolina Duque Castano - Mateus Gustavo Chuqui - Fabiana da Silva Paula - Renato Gamba Romano - William Soares Gattaz Brandão - Frederico P. Brandini - Vivian H. Pellizari.

## Abstract

**Background:** The pelagic environment may present a mosaic of biogeographical domains that regional oceanographic processes can influence. Here, a coastal-to-open ocean microbiome investigation was conducted on 64 water samples from the Santos Basin (SB), South Atlantic Ocean. Using metagenomics and machine learning approaches, we assessed the diversity and distribution of pelagic microbes, identified key bacterial and archaeal taxa, and inferred their ecosystem functions.

**Results:** Unsupervised machine learning revealed a clear spatial and vertical (light availability) distribution pattern across SB, with some indicator taxa previously observed in other marine waters. Supervised learning further revealed that environmental variables, notably phosphate, salinity, and nitrate, which are key markers of local upwelling and the La Plata River plume, are primary drivers of microbial community structure. Furthermore, we recovered 307 metagenome-assembled genomes with 45% of *Archaea* and 42% of *Bacteria* possible new taxa. In terms of functionality, the SB dataset revealed a pelagic ecosystem resembling typical marine (e.g., Atlantic Ocean) waters, with photoautotrophs and nitrogen fixers in the photic zone and different autotrophic pathways in the aphotic environment. Surprisingly, the SB dataset revealed genes for CO bio-oxidation and algal dimethylsulfoniopropionate (DMSP) degradation at all depths. Furthermore, we observed potential non-cyanobacterial diazotrophs in dark water.

**Conclusions:** Our results revealed that the SB represents a unique ecosystem with local oceanographic processes shaping the distribution of diverse and uncharacterized microbiomes. Additionally, these findings highlight the importance of mixotrophic microbes in SB biogeochemical cycles. This massive investigation of the SB pelagic microbiome provided knowledge-based data for understanding local ecosystem health, services, and dynamics, which are essential for future sustainable ocean management.

## Background

The ocean is the largest biome on Earth [1]. They are heterogeneous in terms of scale depth, nutrient availability, microbial distribution, dispersion, and interaction. These habitat variations in the pelagic environment directly impact the function and ecosystem services of the planet [2]. For example, in the photic layer, photoautotrophic microorganisms are responsible for 46% of global primary productivity, an important fraction of the total atmospheric carbon fixation by photosynthesis [3,4,5]. In both photic and aphotic waters, heterotrophic microorganisms contribute significantly to the remineralization of nutrients, organic matter, and carbon export to trophic levels and the deep ocean [6]. In addition to being important for marine primary productivity, the chemical process involves dark carbon fixation via the oxidation of inorganic compounds (e.g., ammonium and sulfur) [7].

The metabolic diversity of prokaryotes in the water column plays an important role in marine biogeochemical cycles [8,9]. Over the past 10 years, circumnavigation expeditions combined with omics advances have significantly improved the study of microbial metabolic functions and their contributions to these cycles [10,11,12,13,14,15]. Despite the increasing number of studies on the global ocean microbiome, knowledge of South Atlantic Ocean (SAO) planktonic prokaryotic metabolism remains limited [16,17]. Most studies on the SAO microbiome have focused on prokaryotic abundance, carbon biomass [18,19,20], and taxa diversity [21,22,23,24]. Until recently, genomic data have suggested that the SAO microbiome has poorly characterized *Bacteria* and *Archaea* taxa [16,17]. This evidence also revealed small- to large-scale changes in the pelagic realm and microbiomes, as well as in the ecosystem services they provide.

The Santos Basin (SB), located in the SAO, is an ecologically and economically important microbial environment. SBs support, regulate, and provide SAO ecosystem services, especially fisheries and fossil energy resources [25]. As a typical deep-water petroliferous region, this basin faces several environmental challenges, including coastal pollution [26,27,28], overfishing [29] and anthropogenic-induced climate change negative effects [30]. Considering that SB is a changing environment under constant threat, characterizing the SB marine microbiome via metagenomic sequencing and machine learning (ML) approaches in a relatively healthy state is crucial [31]. These advanced methods provide an effective way to uncover the complexity of marine microbial communities [32,33,34], offering critical insights essential for understanding, preserving, and managing anthropogenic impacts on marine ecosystems [31,35].

In this study, we investigated the diversity, function, and distribution of planktonic prokaryotic communities in the Santos Basin (South Atlantic Ocean) through the combination of metagenomics and machine learning approaches. Our main goal was to understand (1) the taxonomic and functional diversity of the pelagic bacterial and archaeal communities from different depths across the basin; (2) the recovered genomes and their potential roles in the local biochemical cycle; and (3) the oceanographic features that influence the microbial community. To address these objectives, we collected coastal-to-open ocean water samples and sequenced the metagenomes on an Illumina platform.

## Methods

### Sampling strategy

A total of 67 water samples were collected during the SANAGU 2019 expedition, corresponding to 28 different sampling stations distributed across the Santos Basin (**Fig. 1**). Six discrete sampling depths (1 M, deep chlorophyll maximum layer [DCM], 250, 900 and 2300 M) were chosen within the water masses (Tropical Water, Coastal Water, South Atlantic Central Water, Antarctic intermediate Water, and Upper Circumpolar Deep Water) at each oceanographic station. These depths were defined via the temperature, salinity and fluorimeter profiles. All samples for biological and chemical analysis, in addition to physical data (temperature [°C], salinity [psu], dissolved oxygen [mg/L], colored dissolved organic matter [CDOM, mg.L-¹] and fluorescence [Relative Fluorescence Units, RFU]) were obtained via a combined Sea-Bird CTD/Carrousel 9 Plus system with Niskin bottles and equipped with sensors. For molecular biology, seawater (15 L) was filtered through a peristaltic pump with 0.22 μm membrane pores (Sterivex, Millipore, MA). The filters were stored at −80 °C for 30 days until DNA extraction. Samples for inorganic nutrients were also filtered through 0.22 μm filters onboard and frozen (at −20 °C) until analysis (up to 30 days), with further details from the SB scientific cruises reported in [24].

**Figure 1.**
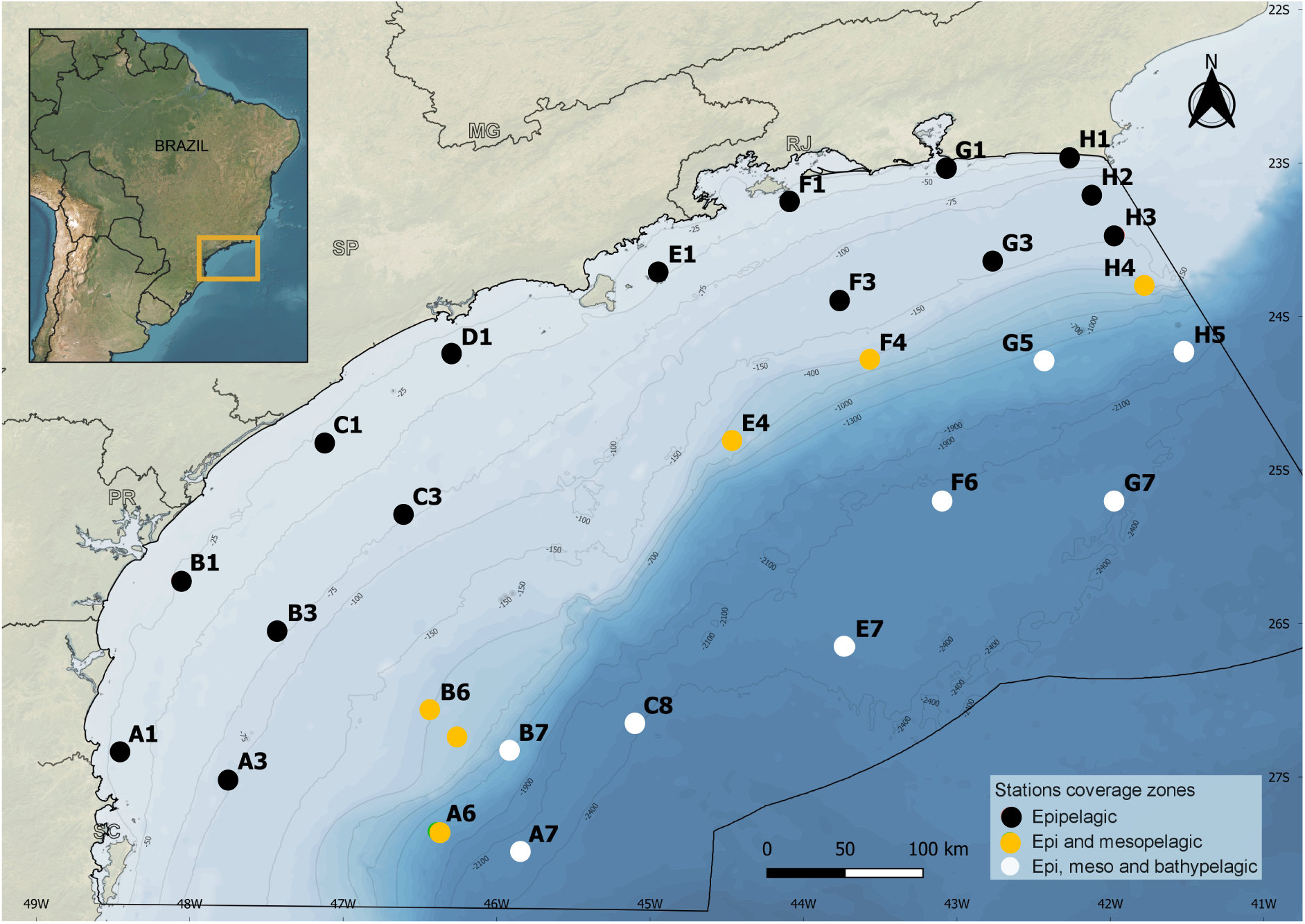
Sampling map of the Santos Basin in the South Atlantic Ocean. Basin localization related to the Brazilian coast and oceanographic stations sampled from August–October 2019. The colored dots represent station coverage sampling zones: black for epipelagic samples; orange for epipelagic and mesopelagic samples; and white for epipelagic, mesopelagic and bathypelagic samples. Abbreviations represent the epipelagic (Epi) and mesopelagic (Meso) zones.

### Environmental factors

Concentrations of inorganic nutrients (nitrate, nitrite, phosphate, and silicate) were determined using a flow injection auto-analyzer (Auto-Analyzer 3, Seal *Inc.*), after filtration through 0.22-μm filters. Ammonium concentration was analyzed in a Hitachi U1100 spectrophotometer. The dissolved organic carbon (DOC) concentration was determined using an Elementar® Vario TOC CUBE. Further details from environmental factor analyzes were reported by [24].

### DNA extraction and sequencing

DNA extraction was performed using the DNeasy PowerWater (Qiagen, USA) according to the manufacturer’s instructions. Negative (no sample) extraction controls were used to ensure the extraction quality. DNA integrity was determined after electrophoresis in a 1% (v/v) agarose gel prepared with TAE 1X (0.04 M Tris, 1 M glacial acetic acid, 50 mM EDTA, pH 8) and staining with SYBR Green (Thermo Fisher Scientific, São Paulo, Brazil). DNA integrity and quantification were determined via a Qubit dsDNA HS assay kit (Thermo Fisher Scientific, São Paulo, Brazil) according to the manufacturer’s instructions.

DNA library construction and sequencing via Illumina technology were conducted at the National Laboratory of Bioinformatics at LNCC (https://www.gov.br/lncc/pt-br, Rio de Janeiro, Brazil). The samples were arranged on sequencing plates on the basis of their DNA concentration. Samples with similar DNA concentration ranges (ng) were grouped on the same plate, as the number of PCR amplification cycles required for library preparation is dependent on the initial DNA quantity, according to the Illumina protocol. In summary, libraries were prepared using the Nextera DNA Flex Library Preparation Kit (Illumina, USA) following the manufacturer’s instructions. Sequencing was performed on the NextSeq 500 System (Illumina, USA) with the NextSeq 500/550 High Output Kit v2.5 (300 cycles), generating 2 × 150 bp paired-end reads.

### Taxonomic, functional inference and machine learning of metagenomic reads

#### Taxonomic and functional inference

First, the raw data were processed via BBduk software (http://jgi.doe.gov/data-and-tools/bb-tools/). Illumina adapters, PhiX and reads with Phred scores below 20 were removed with the following parameters: minlength = 50, mink = 8, qout = auto, hdist = 1 k = 31, trimq = 10, qtrim = rl, ktrim = l, minavgquality = 20 and statscolumns = 5. Prokaryotic taxonomic annotation of short reads was assigned through best-hit classification using Kraken [36,37]. Additionally, functional profiles of short reads were generated through the SEED subsystem and KEGG, which are available at MG-RAST [38,39], with the following parameters: e value of 0.00001, minimum 50 bp alignment and > 60% identity. The top hits were analyzed in search of specific genes and pathways with the SEED database. Hits were normalized by the total number of hit counts for each sample, and a row Z score was calculated to assess significant differences between samples for each metabolic pathway. The functional analyses were performed with R (R Development Core Team) with the packages PhyloSeq [40], ggplot2 [41], and vegan [42].

#### Machine learning

Machine learning was performed with the read’s taxonomic annotation and environmental data (**Table S1).** A hybrid approach that combines unsupervised and supervised learning methods was employed [43] using the iMESC—An Interactive Machine Learning App for Environmental Science [44]. As an unsupervised analysis, we constructed a self-organizing map (SOM) via a neural network method [45] to group similar samples into neurons, i.e., best-matching units (BMUs). In this study, a multilayered SOM approach was utilized. Owing to the high diversity of genera (2.232 genera) identified by the read annotation (**Table S2**), we transformed the genus-level data via the Hellinger method to explore the microbiome distribution in the SB. The first layer incorporated the microbial genus level with a weight of 0.95 on the basis of the Euclidean similarity index. The second layer integrates sample coordinates and depth information (scaled and centered) with a weight of 0.05 and uses Euclidean distance. We followed these steps to determine the number of nodes for the SOM [46]: The initial number of map nodes was determined via the heuristic recommendation 5*√x, where x is the number of samples. Next, we determined the eigenvectors and eigenvalues from the autocorrelation matrix. We then set the ratio between the two sides of the grid to match the ratio between the two largest eigenvalues. Finally, we scaled the side lengths so that their product (xdim * ydim) was as close as possible to the number of map units determined above. This process resulted in the use of 42 neurons.

Each neuron in the final SOM map contains a weighted list of microbial genera, termed a codebook. The codebook generated by the SOM analysis was then subjected to hierarchical clustering (using the Ward method with squared differences) to group similar BMUs and their corresponding samples. The number of clusters formed by hierarchical clustering (HC) was determined with a split moving window analysis, which identifies discontinuities in the relationship between the number of clusters and the within-cluster sum of squares. These neurons serve as descriptors of the microbial communities, and they are referred to as taxonomic associations (Assocs). The Shannon and Chao1 diversity indices of each Assoc were compared via Kruskal–Wallis ANOVA and post hoc Dunn tests. In addition to the SOM and HC analyses, we applied IndVal analysis to evaluate the importance of genus-level features as indicators of specific clusters. Each feature was assigned exclusively to the cluster where its association was strongest, ensuring unique indicators for each cluster. The IndVal statistic was calculated with 999 permutations to assess significance.

Following the unsupervised step, each Assoc was utilized as a response variable in a supervised training phase. This phase aimed to identify the optimal oceanographic variables (**Table S1**) for modeling and predicting the microbial community (Assoc) via machine learning classification algorithms. Initially, the samples were split for validation, ensuring a balanced partition across Assoc. The training dataset (80%) was used for model fitting, whereas the test dataset (20%) served for subsequent evaluation. Random forest classification algorithms were employed with 500 trees, a cross-validation method using 5 folds and 10 repetitions, evaluating up to 10 variables randomly sampled (’mtry’) at each split. The performance of each random forest (RF) model was assessed by standard metrics. This included evaluating accuracy, analyzing confusion matrices to understand predictive performance across different associations, and determining feature importance to identify which oceanographic variables had the greatest influence on SB microbiome associations.

### Metagenome assembly, binning, and genome quality control

We choose co-assembly strategy to increase the throughput and maximize the number of recovered MAGs per depth (1 m, DCM, 250 m, 900 m and 2300 m; **Table S1** to check each co-assembly dataset). Several studies have used this strategy, including the reconstruction of genomes from water columns [47,48]. Reads from all samples were co-assembly via MEGAHIT v1.0.6 [49], and contigs smaller than 1000 bp were removed. Reads for each metagenome were mapped to the co-assemblyco-assembly using Bowtie2 v2.3.4.3 with default parameters [50]. To process the FASTA files, each of the 5 assemblies was processed using the anvi’o v8 [51] workflow, which generated a contig database, identified open reading frames via Prodigal v2.60 [52], identified single-copy *Bacteria* and *Archaea* genes with HMMER v3.2.1 [53], and annotated genes with functions through the NCBI Clusters of Orthologous Groups (COG) [54].

Individual BAM files (mapping files) were profiled with a minimum contig length of 2500 bp. Genome binning was performed by the CONCOCT [55] program with default parameters. We then used the anvi’o interactive interface [51] to refine the bins generated manually by CONCOCT. Bins were quality checked through CheckM v. 1.0.7 [56]. The quality of bins was determined as high-quality draft (>90% complete, <5% contamination), medium-quality draft (>50% complete, <10% contamination), or low-quality draft (<50% complete, <10% contamination) metagenome-assembled genome (MAG), according to the genome quality standards suggested by [57].

MAGs were taxonomically identified based on genome phylogeny using GTDB-Tk [58] and annotated through DRAM (v1.0) [59]. In addition to the DRAM annotations, we used the sulfur cycling database to identify genes for inorganic and organic sulfur cycling [60]. The list of 109 genes related with biogeochemical cycles searched in the Santos Basin reconstructed genomes, is shown in detail in **Table S7**. Genomic analyses were performed with R (R Development Core Team) with with the packages ggplot2 [41], and vegan [42].

### Distribution of metagenome-assembled genomes across the Santos Basin

To investigate the distribution of metagenome-assembled genomes (MAGs) across Santos Basin (SB) depths, the relative abundance data were calculated as the number of reads recruited to a contig divided by the total number of reads recruited across all samples, which were computed using the anvi’o v8 pipeline [61]. First, MAGs were profiled via the “anvi-summarize” function, which provides detailed information about each MAG in the *bin_by_bin* and *bins_across_samples* subdirectories. The *bin_by_bin* subdirectory contained per-bin statistics, including mean coverage and relative abundance across samples, whereas the *bins_across_samples* subdirectory aggregated these data for all MAGs in tab-delimited matrices. The relative abundance of each MAG in different samples was retrieved from the “mean_coverage.txt” and “rel_abundance.txt” files within *bins_across_samples*, ensuring consistency and accuracy in quantifying MAG distributions. For visualization, centered log ratio (CLR) transformation was performed by dividing the relative abundance of each MAG by its geometric mean, followed by the natural logarithm of the result. To ensure comparability, all zero values were replaced with a pseudocount (ε = 1e-6). CLR-transformed values were normalized via min–max scaling to a range of 0--1 for visualization. Each depth was processed independently to highlight depth-specific patterns. The data were visualized on heatmaps.

## Results

### Characterization of Santos Basin metagenomic reads through unsupervised learning

The machine learning analysis identified prokaryotes with ecological relevance by assessing the connections (statistically supported) among read taxonomy in each sample (**Figs. 2** and **3**). The SOM analysis stabilized with a learning rate of approximately 0.0006 for the genus data and 0.06 for the geographical data. The SOM network accounted for 88.70% of the data variance, with a mean topographic error of 0.36. Hierarchical clustering analysis, which uses the elbow method and split moving window to determine the dendrogram cut, identified four taxonomic associations (Assoc) (**Fig. S1**) with vertical and spatial distribution patterns across the SB (**Fig. 2A-E**): (i) In the photic environment (1 m and DCM depths), Assoc 1 (purple group) was represented by coastal epipelagic samples, and Assoc 2 (cian group) was prevalent in the oceanic epipelagic samples; (ii) Assoc 3 (orange group) and 4 (red group) were both prevalent in the aphotic environment, occurring in northern and southern samples, respectively (250, 900 and 2300 m depths).

**Figure 2:**
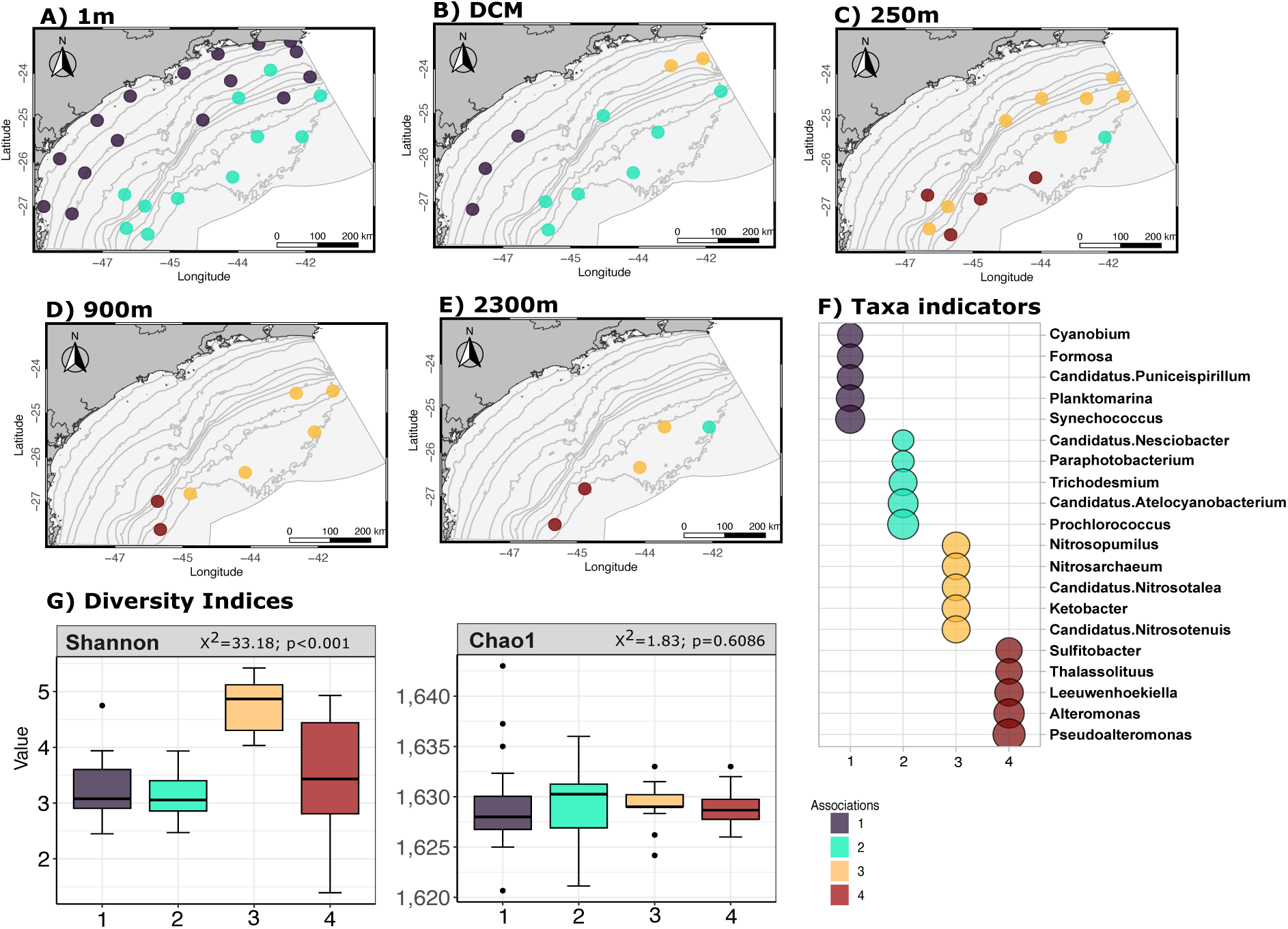
Results from the unsupervised phase of the hybrid model. **A-E:** Spatial distribution of taxonomic associations at each depth across the Santos Basin. **F:** Top 5 significant pelagic-read taxa ecotypes (p < 0.05) for each association, with the point size representing the IndVal statistic, which reflects the strength of the relationship. **G:** Diversity indices (Shannon and Chao1) were calculated for each association, with corresponding chi-square (2) values and p values shown for each comparison. The colors represent the associations: association 1 (black), association 2 (cian), association 3 (orange) and association 4 (red). D) Spatial distribution of associations across depth zones in the Santos Basin.

**Figure 3:**
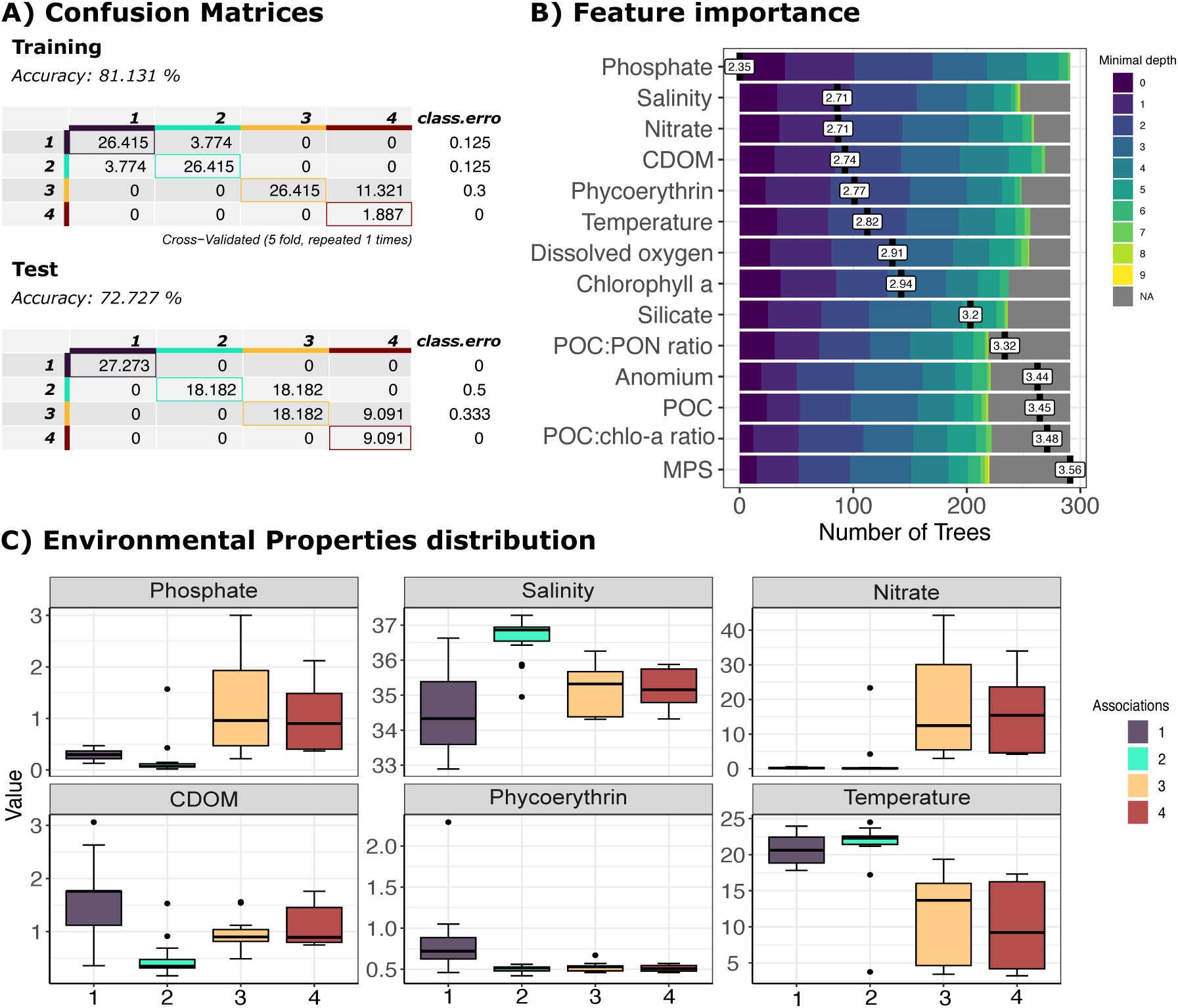
Model performance, variable importance, and environmental properties. **A:** Classification accuracy and errors for training and test datasets, displaying actual versus predicted classes. **B:** Importance of the environmental variables in the random forest model, colored according to the minimal tree depth. **C:** Box plots showing the distributions of key environmental properties across the four associations. The colors represent the associations: association 1 (black), association 2 (cyan), association 3 (orange) and association 4 (red).

The microbial communities across the four taxonomic associations were composed mainly of *Bacteria* (93.2%) rather than *Archaea* (6.7%) (**Table S2**). We focused on the 15 most abundant genera and 5 indicator taxa for each association and observed vertical and spatial distribution patterns (**Fig. S1** and **Fig. 3**). For example, the photic layer (1 m and DCM) generally presented a very similar taxonomic composition, with abundant *Synechococcus* and *Prochlorococcus (*Cyanobacteria) and *Candidatus* Pelagibacter (Assocs 1 and 2, **Fig. S1**). As expected, we detected differences in *Synechococcales* distribution; *Synechococcus* was dominant in the coastal epipelagic samples (Assoc 1), whereas *Prochlorococcus* was dominant in the oceanic waters (Assoc 2, **Fig. S1**). In addition, representatives of *Candidatus* Puniceispirillum*, Formosa, Planktomarina (Roseobacter clade)* and the cyanobacteria *Synechococcus* and *Cyanobium* were indicator taxa for coastal waters, and representatives of *Prochlorococcus*, *Trichodesmium* and *Candidatus* Atelocyanobacterium and the *Bacteria Candidatus* Nesciobacter and *Paraphotobacterium* were indicator taxa for oceanic waters (**Fig. 3E**). The predominant metabolic subsystems of the photic layer (**Fig. S3 and Table S3**) were, as expected, associated with phototrophic lifestyles such as those from *Synechococcales* (photosystems/phycobilisomes) or photoheterotrophy with the rhodopsin pump (generally present in *Candidatus* Pelagibacter).

The taxonomic composition changed as we reached light extinction in the mesopelagic (250 and 900 m depths) and bathypelagic (2300 m depth) zones (Assocs 3 and 4, **Fig. S2**). Spatial distribution differences were observed in the SB dark microbiome. For example, the bacterial representatives *Alternomonas, Pseudoalteromonas* and *Sulfitobacter* were abundant in samples from SB South (Assoc 4), whereas *Candidatos Nitrosomarinus*, *Nitrosopumilus, Gordonia* and *Candidatos Thioblobus* were abundant in samples from the north (Assoc 3). The Archaea *Nitrosopumilus* and *Candidatus* Nitrosopelagicus were prevalent across all aphotic SB (Assoc 3 and 4, **Fig. S2**). In agreement with the dominant taxa, *Nitrosopumilus, Nitrososphaera* and *Candidatus* Nitrosotalea were indicator taxa for the northern region samples, whereas northern region samples indicators were the *Sulfitobacter, Pseudoalteromona* and *Alteromonas,* (**Fig. 3E**).

The aphotic zone of the SB presented a metabolically diverse set of pathways. The highest abundance pathways were associated with nitrate/nitrite ammonification, denitrification (nitrogen gas as the final product) and dissimilatory nitrate reduction (with ammonium as the final product). Other metabolic pathways that coexist in dark water include methanogenesis from methylated compounds, inorganic sulfur assimilation, lysine fermentation and anaerobic respiratory reductases (**Fig. S3** and **Table S3**).

In agreement with spatial taxon distribution and potential metabolic function, significant differences were observed in the Shannon diversity index (Kruskal‒Wallis, χ^2^=33.18, p<0.001) across the four associations (**Fig. 2G**). Post-hoc Dunn tests for the Shannon index revealed that Assoc 3 and 4 had significantly higher median values than Assoc 1 and 2 (Dunn test, p<0.01; **Table S4**). No significant differences were found for the Chao1 richness index (Kruskal‒Wallis, χ^2^=1.88, p=0.6) (**Fig. 2G**). These results suggest that microbial diversity is more pronounced in aphotic environments, as represented by Assocs 3 and 4, whereas richness (Chao1) remains relatively consistent across all the taxonomic associations.

### Predictive modeling of Santos Basin metagenomic reads via supervised learning

We further explored how read’s taxonomy is linked to oceanographic variables in the SB. For this purpose, we combined taxonomic associations (cited above) and environmental data (**Table S1**) through random forest (RF). RF showed an accuracy of 82% for the training dataset and 90% for the test dataset (**Fig. 3A**). Associations 1 and 2 had the lowest error (15%), and association 4 had the highest error (23%) for the training data, whereas association 3 misclassified 50% of the data as association 1 in the test data. Among the 30 environmental variables, 15 were significant (significance level = 0.05). Phosphate was the most important feature, with a mean minimal depth of 2.24, followed by salinity, nitrate, CDOM, phycoerythrin, temperature, dissolved oxygen, chlorophyll-*a* (chlo-*a*), silicate, the POC:PON ratio, ammonium, the POC:chlo-*a* ratio and SPM (**Fig. 3B**). The environmental properties varied significantly across associations (**Fig. 3C**): Assoc 1 presented higher CDOM and phycoerythrin concentrations and moderate salinity and temperature; Assoc 2 presented elevated salinity and temperature levels and lower CDOM; Assoc 1 and Assoc 2 showed elevated phosphate and nitrate concentrations than the other Assocs; Assoc 3 and Assoc 4 presented higher phosphate and nitrate concentrations and salinities and temperatures than Assocs 1 and 2; and CDOM distinguished Assoc 4.

### Genome-recovery metagenomics reveals taxonomic novelty

The co-assembly of pelagic metagenomes, combined with automated binning and manual refinement, allowed the reconstruction of 307 medium- or high-quality Santos Basin metagenome-assembled genomes (SB-MAGs). Among these MAGs, 59 were considered high-quality drafts, and 248 were classified as medium-quality drafts, with an average genome completeness of 76.9%, according to the genome quality standards suggested by Bowers *et al.* (2017). The statistics of the SB-MAGs are described in **Table S5**.

A total of 256 bacterial SB-MAGs were taxonomically assigned to 17 phyla, and 51 archaeal SB-MAGs were assigned to 2 phyla based on the GTDB taxonomy (**Table S4**). *Proteobacteria* (class *Gammaproteobacteria* and *Alphaproteobacteria*, n = 81) followed by *Planctomycetota* (n = 48) were the most abundant bacterial phyla in the SB-MAGs, whereas archaeal MAGs were assigned to the phyla *Thermoplasmatota* (class *Nitrososphaeria*, n = 31) and *Thermoproteota* (class *Poseidonia*, n = 18). The SB-MAGs possibly included taxonomic novelty, with 45% and 42% novel species within the bacterial and archaeal MAGs, respectively (**Fig. 4**). In addition, two genomes recovered from depths of 900 m were not classified by the GTDB.

**Figure 4:**
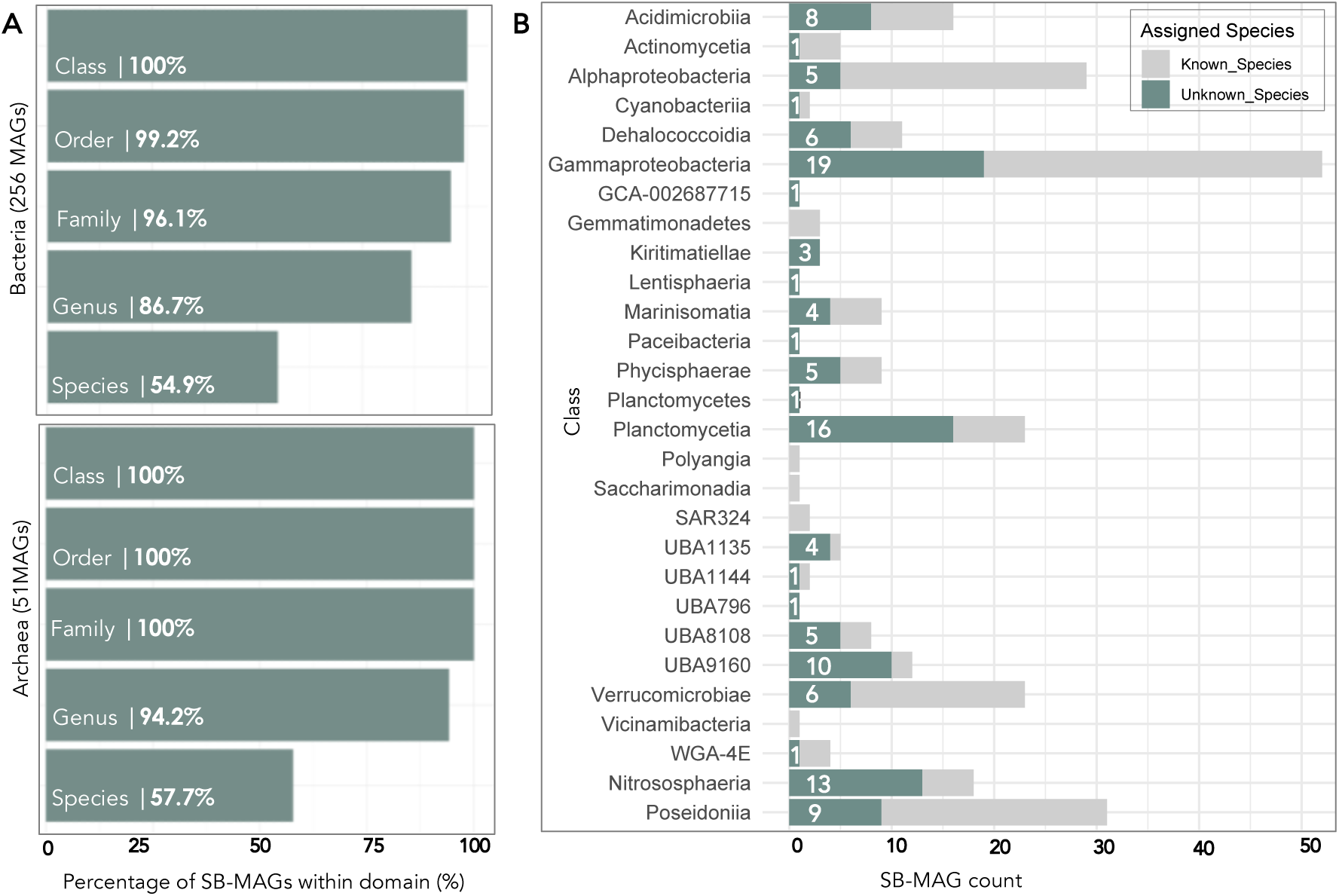
Stacked bar plot for taxonomic novelty of the Santos Basin Metagenome-Assembled Genomes. **A:** Percentage of genomes (X axis) with classification according to their taxonomic ranks (Y axis) for archaea and bacteria. **B:** Number of genomes (X axis) at the class level (Y axis). The taxonomically unclassified portion is depicted in green and classified in gray.

### Genome-resolved metabolic profiles reveal distinct roles in biogeochemical cycles

In general, we recovered procaryotes SB-MAGs from all depths (**Fig. 5** and **Fig. S4**). The number of recovered MAGs increased with depth (from 2 MAGs per 1 m depth co-assembled samples to 10 MAGs per 2300 m depth co-assembled samples), especially at 250 m depth (**Fig. S4**). The MAGs classified as *Alphaproteobacteria* (73% average genome completeness - AGC), *Gammaproteobacteria* (84% AGC) and *Acidimicrobiia* (81% AGC) bacterial classes and the *Nitrososphaeria* archaeal class (79% AGC) encoded the greatest number of genomes with high-level completeness and biochemical cycle metabolic marker genes (**Fig. 5A**).

**Figure 5:**
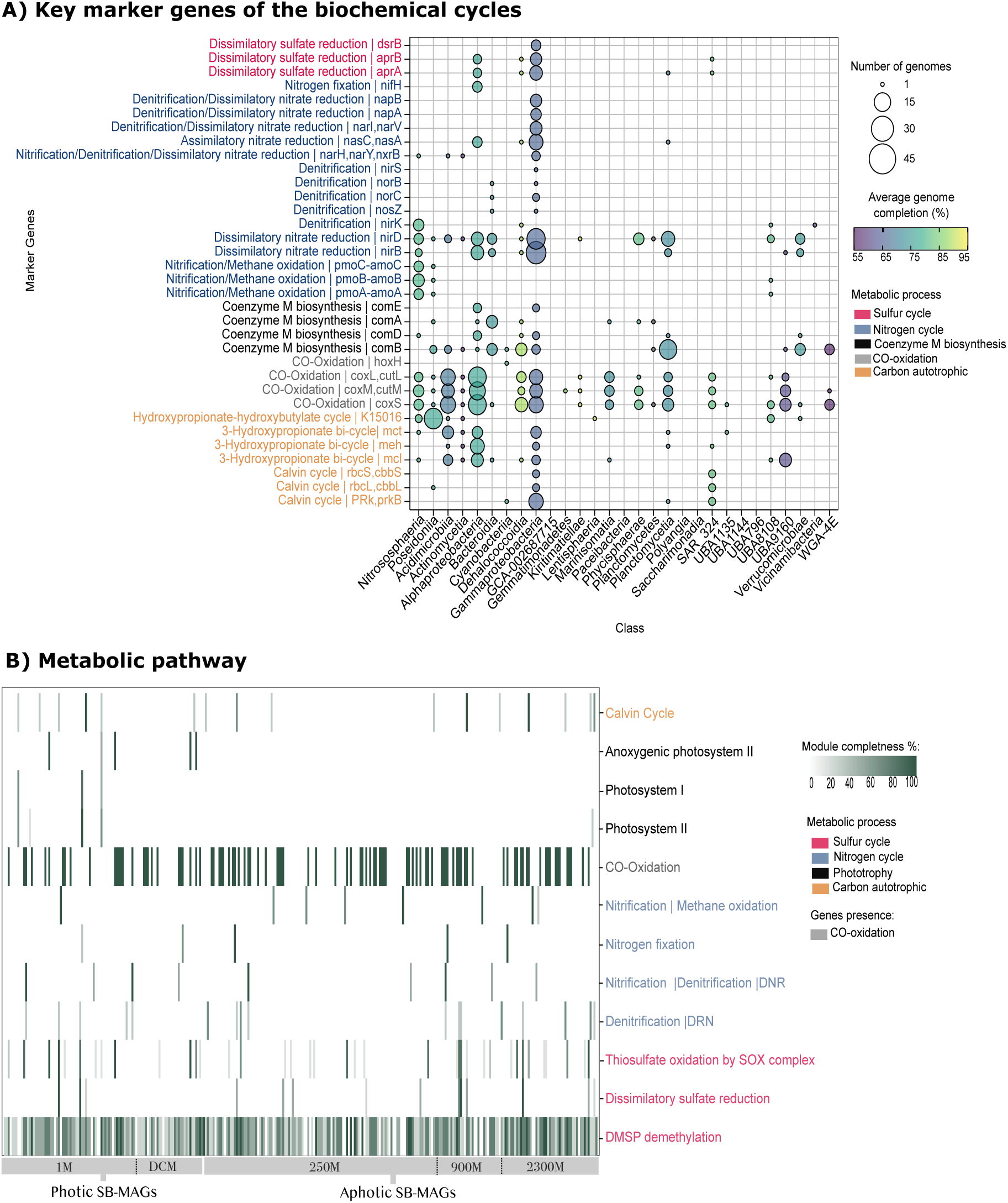
Metabolic potential of 307 Santos Basin metagenome-assembled genomes (SB-MAGs). **A:** Bubble plot showing key marker genes of the biochemical cycles in the SB-MAGs. MAGs are summarized at the class level, with the size of the circle corresponding to the number of genomes in that class with a given gene and the color reflecting the percentage of genome completeness. Marker genes that were not detected in any MAG were omitted. **B:** Heatmap of the percentage of metabolic pathways enriched with the SB-MAGs. The percentage module completeness for each pathway is coded by color intensity. Only module completeness equal to or greater than 60% was plotted.

Among all the MAGs retrieved from this work, 48 MAGs with 60% or more complete metabolic pathways in all the metagenomes were selected, and their main distinguishing metabolic feature genes were identified (**Fig. 5B**). The rest of the MAGs in all the datasets are shown in detail in **Table S7**.

#### MAGs from the photic layer

As expected, key players in photic waters harboring phototrophic/photoheterotrophic lifestyles, such as *Cyanobacteria* and *Rhodobacteraceae* (genus MED-G52), were detected only in the photic datasets (**Fig. 5B**). Subunits from Photosystems I (*psaABCDEF*) and II (*psbABCDEF*) were affiliated with the cyanobacterium *NIES-981 genus* (1M_MAG_14, 91% genome completeness - GC) and *Atelocyanobacterium thalassa_A* spp. (1M_MAG_116, 51% GC, ANI 99%), with 67% and 100% photosynthetic metabolic pathways, respectively (**Fig. 5B**). The *nifK* gene for nitrogen fixation was also detected in the *Cyanobacteria* cited above.

The photoheterotrophic anoxygenic photosynthesis type I reaction center was absent in the MAGs. Photoheterotrophic lifestyle, using all genes (*pufML*) that encoding subunits in the anoxygenic photosynthesis type II reaction center was detected in four *Rhodobacteraceae* MAGs: two in MED- G52 sp002457055 (1M_MAG_46 and PMC_MAG_87, 73% AGC, ANI > 96%) and two in MED-G52 sp001627375 (1M_MAG_193 and PMC_MAG_74, 59% AGC, ANI > 98%) (**Fig. 5B**). These genomes also have complete or 65% thiosulfate oxidation by the SOX complex (*soxABCDXYZ* genes) and marker genes for CO oxidation (*coxL, cutL, coxM, cutM and coxS*) and the 3-hydroxypropionate bicycle (*mct* and *meh*), providing some evidence for chemoautotrophy (the coupling of carbon fixation and reduced compound oxidation). Except for MED-G52 sp001627375 (PMC_MAG_74), no *mct* or *meh* genes were detected.

A family that provided abundant MAGs in SB photic waters was *Pirellulaceae* (10 MAGs with 86% AGC). The exclusive photic Pirellulaceae genomes (ANI > 96%) have genes (*fwdA, fmdA_fwdB, fmdB_fwdC, fmdC* and *ftr*) for 80% of C1 metabolism: *UBA721 sp002698965* (PMC_MAG_07), *UBA721 sp002701385* (1M_MAG_12 and PMC_MAG_01), *GCA-2723275 sp002723275* (1M_MAG_54 and PMC_MAG_08), and *UBA11883 sp002714565* (PMC_MAG_11).

Surprisingly, two genomes of the *UBA868* family (84% of AGC, ANI <97%) have genes for sulfur metabolism: *UBA868 sp016776875* (1M_MAG_66) and *UBA868 sp010025385* (1M_MAG_104).

These genomes have complete or 83% thiosulfate oxidation by the SOX complex and complete dissimilatory reduction of sulfate (*sat* and *aprAB* genes). In addition, *UBA868 sp010025385* has a complete DMSP demethylation pathway (*dmdA, dmdB, dmdC* and *dmdD* genes).

Potential chemoautotrophic metabolism under photic SB was observed in two MAGs: *Novosphingobium indicum* and *Gordonia sp002700145*. *Novosphingobium indicum* (PMC_MAG_59, xx% of AG, ANI <98%) has genes for CO oxidation and nitrogen fixation (*nifH, nifD* and *nifK*) and partial (>70%) DMSP demethylation, a potential non-cyanobacterial diazotroph. *Gordonia sp002700145* (1M_MAG_21 and PMC_MAG_02, 95% AGC, ANI>98%) is potentially chemoautotrophic by reduced nitrogen, with all the genes for CO oxidation and nitrate reduction (*narG, narZ, nxrA, narH, narY* and *nxrB*) and partial DMSP demethylation (>60%).

#### MAGs from twilight and aphotic waters

Some functional traits cooccur in both SB photic and aphotic waters with potential sulfur, CO, and nitrogen metabolism (**Fig. 5B)**. However, these metabolic pathways are prevalent in the twilight and aphotic waters of SB. For example, complete or partial (>70%) DMSP demethylation was identified in various genomes (115/307): *Gammaproteobacteria* (35/307), *Alphaproteobacteria* (22/307), *Acidimicrobiia* (14/307), *UBA9160* (8/307), and *Dehalococcoidia* (7/307). This metabolism was higher in exclusively aphotic genomes than in photic genomes (74/307 vs. 41/307, respectively) (**Fig. 6**). For example, six MAGs have all DMSP demethylation and CO oxidation marker genes: (i) two *Pseudohongiellaceae* families (MAGs with 84% AGC); (ii) *UBA9145 sp002712055* (250M_MAG_01, 90% GC, ANI99%), *UBA9145 sp002694855* (900M_MAG_11, 78% GC, ANI99%), *GCA-002712965 sp009391195* (250M_MAG_89, 81% GC, ANI99%); and (iii) the new UBA9410 (250M_MAG_120, 71% GC, ANI89%) species. Additionally, SAR324 genomes represented by *Arctic96AD-7 sp002082305* (900M_MAG_17 and 2300M_MAG_22, 84% of AGC, ANI <98%) have genes for partial (75%) DMSP demethylation and all genes for the Calvin cycle (P*RK, prkB, rbcL* and *rbcS* genes) and for CO oxidation.

**Figure 6:**
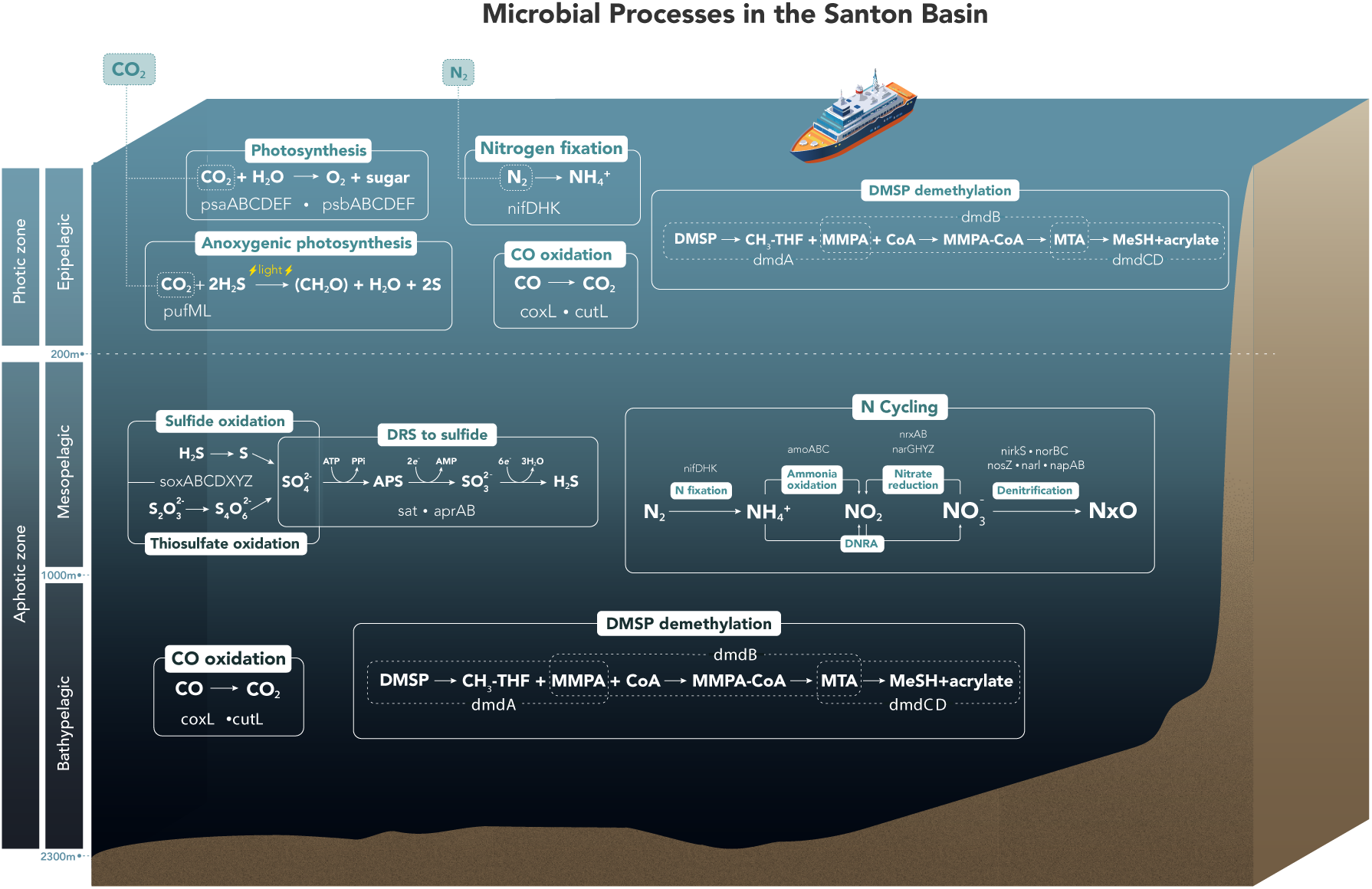
Conceptual diagram of biogeochemical functions in the Santos Basin, South Atlantic Ocean. Metabolic microbial pathways involved in the carbon, nitrogen, and sulfur cycles are shown. Metabolic pathways in the aphotic environment (<200 m depth) were grouped due to the substantial taxonomic and functional redundancies observed in the SB-MAGs. Acronyms: carbon dioxide (CO2), nitrogen gas (N2), ammonium (NH4^+^), nitrite (NO2^-^), nitrate (NO3^-^), nitrogen (N), ammonia (NH4^+^), hydrogen sulfide (H2S), sulfate (SO), thiosulfate (S2O3^2−^), tetrathionate (S4O6^2−^), dimethylsulfoniopropionate (DMSP), methyltetrahydrofolate (CH3-THF), 3-methiolpropionate (MMPA), coenzyme A (CoA), 3- methiolpropionyl-CoA (MMPA-CoA), methanethiol (MTA or MeSH), adenosine triphosphate (ATP), pyrophosphate (PPI), adenosine monophosphate (AMP), and adenosine 5’-phosphosulfate (APS).

Another abundant form of sulfur metabolism involves thiosulfate oxidation by the SOX complex (39/307 MAGs) (**Fig. 5A**). Complete or partial (60–80%) SOX complex (s*oxABCDXYZ*) marker genes were found in 17 bacterial genomes and were more abundant at deep-sea MAGs than at surface MAGs (10/307 vs. 7/307, respectively) (**Fig. 5B**). A partial Sox system (60–80%) and DMSP demethylation (<75% CMP) were detected in 2 MAGs represented by *Sulfitobacter pontiacus* (250M_MAG_190 and 2300M_MAG_15, 69% AGC, ANI 97%) and possible new *Halioglobus_A* (250M_MAG_26, 94% GC) species. The 2300M_MAG_15 and 250M_MAG_117 genomes also have all CO oxidation genes.

The microbial sat and *aprAB* genes encoding enzymes involved in dissimilatory reduction of sulfate (DRS) to sulfite were detected in teen MAGs (**Fig. 5B**). However, complete or 75% of the marker genes *aprAB* and *drsAB* for the DRS pathway were limited to three *UBA868* family (MAGs with 81% AGC, ANI <97%) genomes: *REDSEA-S09-B13 sp002456995* (900M_MAG_12) and *UBA11791 sp0027255* (900M_MAG_14 and 2300M_MAG_19) (**Fig. 5B**). These genomes also have genes for partial (83%) thiosulfate oxidation and all genes for CO oxidation. Additionally, 900M_MAG_12 and 2300M_MAG_19 have partial (75%) DMSP demethylation.

The exclusive deep-sea genomes (4 MAGs) with chemoautotrophy by sulfur metabolism were as follows: (i) the *Casp*-*alpha2* family (MAGs with 60% AGC, ANI <98), *Casp-alpha2 sp002938315* (2300M_MAG_52) and *Casp-alpha2 sp002938075* (2300M_MAG_70), which included complete DMSP demethylation and all genes for CO oxidation. Additionally, *Casp-alpha2 sp002938075* has a complete Sox system and a partial DRS pathway (50%), and *Casp-alpha2 sp002938315* has a partial Sox system (>60%); (ii) new *Pseudohoeflea* (250M_MAG_117, 65% of GC) species with complete DMSP demethylation, a partial Sox system (>60%) and all genes for CO oxidation; and (iii) *Pelagibacter* (2300M_MAG_37, 83% of GC) with partial (>60%) DMSP demethylation genes for CO oxidation and urea metabolism (*UreB* and *UreC* genes).

Next, we identified the main nitrogen ecological drivers, which included potential exclusively ammonia-oxidizing archaeal *Nitrososphaeria* (14/307) and non-cyanobacterial diazotroph genomes (03/307). The ammonia-oxidizing archaea *Nitrosopelagicus brevis* (3 MAGs, 250M_MAG_84, 900M_MAG_29 and 2300M_MAG_25, 61% of AGC, ANI >96%), *Nitrosopumilus sp002690535* (250M_MAG_61, 75% of GC, ANI 95%), and two possible novel species, *Nitrosopelagicus* (250M_MAG_164, 56% of GC) and *Nitrosopumilus* (250M_MAG_116, 67% of GC), have more than 60% of the genes required for complete/partial nitrification (*amoABC* genes). In all these cited MAGs, we detected the key enzyme for the archaeal 3-hydroxypropionate–4-hydroxybutyrate autotrophic carbon dioxide assimilation pathway (K14534: abfD; 4-hydroxybutyryl-CoA dehydratase/vinylacetyl- CoA-delta-isomerase, except for the 250M_MAG_164 and 900M_MAG_29. The potential ammonia- oxidizing *bacteria MarineAlpha9-Bin7 sp002937585* (2300M_MAG_54, 64% of GC, ANI 99%) have all of the genes required for CO oxidation and the key gene for nitrification (*hao*). Surprisingly, members of the Sphingomonadaceae family have all genes related to nitrogen fixation (*nifH, nifD* and *nifK*) and CO oxidation and partial (>70%) DMSP demethylation with a unique taxonomic distribution: a potential non-cyanobacterial diazotroph *Novosphingobium indicum* (250M_MAG_22, 900M_MAG_06 and 2300M_MAG_05, 91% AGC, ANI <98%).

The *NarG, narZ, nxrA, narH, narY* and *nxrB* genes involved in nitrate reduction were detected in two Gordonia MAGs: *Gordonia sp002700145* (250M_MAG_32 and 900M_MAG_05, 94% AGC, ANI>88%). These *Gordonia* MAGs also included all genes involved in CO oxidation and partial DMSP demethylation (>60%), with potential chemoautotrophic effects that reduce nitrogen.

Marker genes for denitrification (*nar, nir, nor* and *nos*) were detected in seventeen SB-MAGs (**Fig. 5A**). Only four *Pseudomonadales* and one *Enterobacterales* novel species had more than 60% complete/partial denitrification: two *Ketobacter* genera (250M_MAG_05 and 2300M_MAG_02, 98% AGC), one *Halioglobus* genus (250M_MAG_26, 94% GC), one *Oleispira* genus (2300M_MAG_50, 97% GC), and one *Alteromonadaceae* (2300M_MAG_71, 57% GC) (**Fig. 5B and Table S7**).

The dissimilatory nitrate reduction to ammonia (*DNRA, nirBD* or *nrfAH* genes) was a widespread pathway in the pelagic SB (70/307) (**Fig. 5A**). However, no MAG represented more than 60% of the complete DNRA pathway.

An important ecological family in SB waters was *Alteromonadaceae* (8 MAGs with 86% AGC), with more genomes recovered from deep-sea MAGs than from surface MAGs (6/307 vs. 2/307, respectively) (**Fig. 5A**). The genomes were related at the GTDB species level (ANI > 97%) to *Alteromonas macleodii* (4 MAGs), *Alteromonas mediterranea* (1 MAG), *Pseudoalteromonas arabiensis* (2 MAGs), and one possible new *Alteromonas* species (250M_MAG_55). Finally, genes for CO oxidation were predominant in the SB (99/307 MAGs), as described above (**Fig. 5A**). All the key microbial processes associated with SB are summarized in **Fig. 6**.

### The relative abundance of Santos Basin genomes aligns with machine learning results

Despite the co-assembly strategy, the differences in the relative abundances of the 48 genomes explored in the previous section aligned with the SOM association distribution patterns at each depth (**Fig. S5** and **Fig. S6**). For example, in the photic layer, in the MAGs recovered from a depth of 1 m, high relative abundances of the *UBA86* clade (1M_MAG_66 and 1M_MAG_104), the cyanobacterium *NIES-981 genus* (1M_MAG_14) and MED-G52 sp001627375 (1M_MAG_193) were observed in coastal waters (Assoc 1), whereas *Pirellulaceae* MAGs (1M_MAG_12 and 1M_MAG_54) and the cyanobacterium *Atelocyanobacterium thalassa_A* spp. (1M_MAG_116) were enriched in oceanic waters (Assoc 2). Some genomes presented relatively high abundances in both coastal and oceanic photic samples (Assocs 1 and 2): *Gordonia sp002700145* (1M_MAG_21) and MED-G52 sp002457055 (1M_MAG_46). Within the genomes from the DCM layer, four *Pirellulaceae* MAGs (PMC_MAG_01, _07, _08 and _11) were enriched in the oceanic waters (Assoc 2), whereas the *Rhodobacteraceae* MAG (PMC_MAG_87) was enriched in the oceanic waters (Assoc 2). Some genomes presented a higher relative abundance of 2 associations: the *Rhodobacteraceae* MAG (PMC_MAG_74) in both coastal photic samples (Assoc 1) and north aphotic samples (Assoc 3) and the *Sphingomonadaceae* MAG (PMC_MAG_59) in both oceanic photic (Assoc 2) and north aphotic (Assoc 3) samples (**Fig. S5**).

In the aphotic waters, some MAGs recovered from the 250 m depth, such as the Ketobacteraceae (250_MAG_05), *Sphingomonadaceae* (250_MAG_22), *Pseudomonadales* (250_MAG_26), *Nitrosopumilaceae* MAGs (250_MAG_61 and 250_MAG_116) and *Rhizobiaceae* (250_MAG_117) families, presented relatively high abundances in the northern aphotic samples (Assoc 3), whereas the *Alteromonadaceae* (250_MAG_55), UBA826 (250_MAG_89) and *Rhodobacteraceae* (250_MAG_190) MAGs were enriched in the southern aphotic samples (Assoc 4). Compared with those of other associations, the genomes of *Mycobacteriaceae* (250_MAG_32) and *Nitrosopumilaceae* MAGs (250_MAG_84 and 250_MAG_164) were enriched in oceanic waters (Assoc 2). Two genomes showed a higher relative abundance in both northern and southern aphotic samples (Assoc 3 and 4): *Pseudohongiellaceae* (250_MAG_01) and *Acidimicrobiales* (250_MAG_120). At the 900 m depth, the *Mycobacteriaceae* MAG (900M_MAG_05) had a high relative abundance in the northern aphotic samples (Assoc 3), whereas the *Nitrosopumilaceae* MAG (900M_MAG_29) was enriched in the southern aphotic samples (Assoc 4). Three genomes, *Sphingomonadaceae* (900M_MAG_06), *Pseudohongiellaceae* (900M_MAG_11) and UBA868 (900M_MAG_12), were enriched in both northern and southern aphotic samples (Assoc 3 and 4). Finally, at 2300 m depth (Associations 2 and 4), a greater abundance of *Nitrosopumilaceae* MAG (2300_MAG_37) was observed in the oceanic waters (Assoc 2) than in the other associations (**Fig. S6**). Patterns of MAG distributions at each depth and association highlight potential ecological adaptations to varying environmental conditions. The full SB-MAG abundances across all datasets are detailed in **Table S8**.

## Discussion

This study revealed that the Santos Basin (SB) represents a unique ecosystem harboring diverse microbiome. The key SB prokaryotic taxa and their ecosystem functions differed between the photic and aphotic depths. Essential nutrient cycling (carbon, nitrogen, and sulfur) has the potential to affect SB ecological dynamics and is encoded by uncharacterized taxa at the genus and species levels. This finding provides evidence that microbial taxa in subtropical regions are highly underexplored [33]. The inferred metabolic pathways and related enzymes from the recovered genomes constitute a first step in understanding the role of the SB pelagic microbiome in maintaining ecosystem functions.

### Factors influencing microbial diversity and distribution patterns

The influence of local oceanographic features on the SB microbiome distribution was powerfully predicted via machine learning. The SB coastal waters favored nutrient-rich taxa, whereas the photic oceanic waters supported those suited in an oligotrophic environment. Associations 1 and 2 were found in the photic environment, dominated by photoautotrophic and photoheterotrophic lifestyles adapted to light and nutrient variations. In contrast, Associations 3 and 4 were characteristic of aphotic and deep waters, with nitrogen-fixing and sulfur-oxidizing taxa reflecting adaptation to deeper nutrient-rich waters. The SB microbial associations were influenced by phosphorus nitrate, CDOM, dissolved oxygen, salinity and light availability, resulting in clear spatial and vertical distributions. Salinity and temperature were previously reported as the major drivers of marine microbial community composition in the global ocean [10,13] and South Atlantic Ocean [16,17, 24]. The nutrient concentration was recently predicted to be the most important driver of microbial ecology alterations under low-level oceanic climate change scenarios [62].

Moreover, it is important to emphasize that salinity and temperature are the main features of water masses with specific concentrations of nutrients, dissolved oxygen, and organic matter [63]. Water masses are influenced by different medium-scale oceanographic processes, such as mesoscale eddies, upwelling and mixing. High concentrations of phosphate and nitrate are possibly key markers of two important coastal oceanographic processes in the SB: upwelling of South Atlantic Central Water in the northern portion and the La Plata River plume influence in the southern portion [24,25, 64]. These processes can potentially increase biomass productivity [18, 64], leading to an increase in prokaryotic taxa and associated ecosystem functions. These distinct environmental profiles across the four associations illustrate the complexity of nutrient-driven and physical processes shaping the SB microbiome. This underscores the combined influence of local oceanographic phenomena in structuring microbial distributions.

### Santos Basin microbiome

The first ecogenomic insights into the SB water column revealed that this microbiome has some typical indicator taxa previously observed in other pelagic environments [10,17, 65]. Additionally, SB included cyanobacterial diazotrophs as indicator taxa. For example, the cyanobacteria *Synechococcus,* heterotrophic *Planktomarina* and nitrogen-fixing *Trichodesmium* were dominant in coastal photic SB waters. Conversely, the photoautotrophs *Prochlorococcus* and *the* nitrogen-fixing *Candidatus* Atelocyanobacterium (UCYN-A) are prevalent in oceanic photic waters. Although *Synechococcales* genomes from SB were not recovered, photoautotrophic and nitrogen fixation activities were confirmed through gene recovery *NIES-981* and *Atelocyanobacterium thalassa A* (UCYN-A). The most abundant *Prochlorococcus’* sympatric photoheterotrophic is the *Pelagibacterales* clade (SAR11) [66,67]. This clade was abundant in all SB photic waters, not only oligotrophic oceanic waters, contributing to SB dissolved organic matter remineralization.

The potential prevalence of carbon monoxide (CO) bio-oxidation in the SB pelagic environment supports recent studies in which the *coxL* gene was widely distributed in photic [68] and dark waters [13]. In fact, SB waters presented a higher diversity of CO metabolizers. These microorganisms may use dissolved CO as a carbon and energy source [13,69, 70,71,72]. For example, in SB photic waters, potential aerobic anoxygenic phototrophs (AAPs) could use light and CO oxidation as complementary energy sources, as previously reported [73,74]. Genes for aerobic anoxygenic photosynthesis (*pufLM*) and CO oxidation were present in MED-G52 (order *Rhodobacterales*). AAPs harness light energy to accumulate organic carbon, which would otherwise be respired, and use it for growth [75]. All AAP taxa recovered in the SB also carried genes for thiosulphate oxidation, providing evidence that these taxa use sulfur as an electron donor for photosynthesis, as previously observed [33,76].

Similar to previous works [77,78], SB exhibited increased biodiversity in deep-sea ecosystems (<200 m depth). This could be related to the higher recovery of MAGs at this depth, especially among mixotrophic and chemolithotrophic CO oxidizers. Although the ecological implications of mixotrophy for marine ecosystem dynamics are unclear [79], it has been suggested to increase dissolved inorganic carbon (DIC) fixation and increase the efficiency of the biological carbon pump [80,81]. In the SB dataset, SAR324 and *UBA868* emerged as potential mixotrophic groups, as previously described [7, 66]. Notably, only SAR324 possesses genes for DIC fixation by *RuBisCO* genes, whereas both SAR324 and *UBA868* harbor genes for degrading dissolved organic sulfur compounds, including dimethylsulfoniopropionate (DMSP), underscoring their roles in carbon and sulfur cycling.

The ecological importance of chemolithoautotrophs in the dark ocean is well established [13, 82,83,84], although our knowledge of the key microorganisms driving this process in deep waters remains unclear [13]. In the aphotic, oxygenated, and nutrient-rich deep waters of SB, *bacteria* and *Archaea* use DMSP (organic compound), thiosulfate/sulfide and ammonia (reduced inorganic compounds) as energy sources for dark carbon fixation. Chemolithoautotrophs in the deep SB were suggested by the presence of the ammonia oxidizers *MarineAlpha9-Bin7 sp002937585, Nitrosopelagicus brevis* and *Nitrosopumilus sp002690535*. Another potential SB chemolithotroph by CO and thiosulfate bio-oxidation is Casp-alpha2, as previously reported in other regions [85]. The *Casp*-*alpha2* genomes of SB also carried genes for degrading DMPS, indicating a potential relationship with the degradation of dissolved organic sulfur.

Surprisingly, potential non-cyanobacterial diazotrophs, *Novosphingobium indicum*, were observed in the deep SB. The ecological importance of these deep-sea diazotrophs is unclear because of the availability of carbon substrates to support their energetic demand for N2 fixation [13,86]. *Novosphingobium* members are known to be metabolically versatile and are usually associated with the biodegradation of aromatic compounds [87]. Interestingly, in addition to the *nif* genes, these *Novosphingobium* genomes have genes for DMSP degradation and CO oxidation, indicating potential deep-sea chemolithoautotrophic diazotrophy.

Recent research has highlighted an increase in sulfur oxidation activity in the deep Atlantic Ocean [17,88]. In fact, the SB dataset reveals significant potential for the degradation of dissolved organic sulfur compounds across the pelagic realm, with a notable focus on alga-derived dimethylsulfoniopropionate (DMSP). This compound plays a dual role in marine systems: serving as a critical sulfur and carbon source for microbial communities and acting as a defense mechanism for phytoplankton [89]. Among the ecologically relevant taxa involved in the sulfur cycle in the deep SB, *Sulfitobacter pontiacus* was identified as an ecologically relevant heterotrophic bacterium. Genes for the DMSP demethylation pathway were found in these SB genomes, which is consistent with the role of *Sulfitobacter* in DMSP degradation in marine environments [90,91,92].

The ecological roles and ecosystem services of the pelagic microbiome have conventionally been associated with the activities of autotrophs (phototrophs or chemoautotrophs) and heterotrophs. Consequently, the organic matter sources supporting deep ocean microbial metabolism come from the euphotic layer (biological carbon pump) or are generated by chemoautotrophic ammonia oxidizers [93]. In the SB, we confirmed the presence of autotrophs and heterotrophs in the pelagic realm. Additionally, this study reveals potential mixotrophic activity in the biogeochemical transformations of organic sulfur compounds in the deep SB and highlights the importance of mixotrophs in South Atlantic Ocean biogeochemical cycles.

## Conclusions

Our results revealed that (1) the prokaryotic taxonomic and diversity in the SB differed between photic coastal and oceanic waters and aphotic northern and southern waters, suggesting that pelagic microbes reflect differences in regional hydrography and biogeochemistry; (2) the vast array of unclassified genomes in this study confirmed that the Atlantic Ocean microbiome and its function are poorly characterized; (3) metabolic versatility, such as autotrophy (photoautotrophic and chemosynthetic), heterotrophy, mixotrophy (chemosynthetic and heterotrophic) and diazotrophy, highlights adaptive strategies that contribute to SB microbiome resilience; (4) genome distribution patterns at each investigated depth and association, suggesting potential niche specialization with ecological adaptations to biotic/abiotic factors; and (5) the SB microbiome may be sustained by dissolved nitrogen compounds (ammonia) and by sinking particles containing organic carbon and sulfur as energy sources, which are supplied by the photosynthetic ecosystem at the ocean surface.

Overall, this study enhances our understanding of Santos Basin microbial ecology, reveals complex relationships among regional oceanographic processes, and emphasizes the use of machine learning in microbial oceanography. These insights are crucial for understanding how biological communities may respond to anthropogenic activities and climate change.

## List of abbreviations

Santos Basin: SB
Dimethylsulfoniopropionate: DMSP
South Atlantic Ocean: SAO
Machine learning: ML
Deep chlorophyll maximum layer: DCM
Colored dissolved organic matter: CDOM
Chlorophyll-*a*: chlo-*a*
Suspended particulate matter: SPM
Particulate organic carbon: POC
Particulate organic nitrogen: PON
Dissolved organic carbon: DOC
An Interactive Machine Learning App for Environmental Science: IMESC
Self-organizing map: SOM
Best-Matching Units: BMUS
Taxonomic associations: Assoc Hierarchical clustering HC
Random forest: RF
Metagenome-assembled genome: MAG
Kyoto Encyclopedia of Genes and Genomes Orthology: KOS
Centered log ratio: CLR
Santos Basin metagenome-assembled genomes: SB
MAGs Average genome completeness: AGC
Genome completeness: GC
Carbon monoxide: CO
Aerobic anoxygenic phototroph: AAP
Dissolved inorganic carbon: DIC
Dimethylsulfoniopropionate: DMSP

## Declarations

### Ethics approval and consent to participate

Not applicable.

### Consent for publication

All the authors have read and approved the paper for submission.

### Availability of data and material

The sequencing read dataset generated and analyzed during the current study is available from the National Center for Biotechnology Information (NCBI) repositor under BioProject PRJNA1191028. The genomic dataset generated during and analyzed during the current study is available in the Figshare repositor under DOI: https://doi.org/10.6084/m9.figshare.27651369.v1.

### Competing interests

The authors declare that they have no competing interests.

### Funding

This study was funded by Petróleo Brasileiro S.A. (PETROBRAS), through the RD&I investments clauses of the Brazilian National Agency of Petroleum, Natural Gas, and Biofuels (ANP), under grant numbers 5850.0109317.18.9 and 21167-2, respectively. The project “Caracterização química e biológica do sistema pelágico da Bacia de Santos/PCR-BS” is a national collaboration between PETROBRAS and Universidade de São Paulo. NMB and LNL were financed by a Pos Postdoctoral fellowship from the PETROBRAS. ATRV is supported by FAPERJ (E-26/201.046/2022) and CNPq (307145/2021-2).

### Authors’ contributions

N.M.B., V.H.P., D.M. and C.R.J. conceived and designed the study. J.C.F.M., A.M.A., A.M.E., R.G.R., D.C.D.C., F.S.P. and W.S.G.B. was responsible for onboard sampling and extracting DNA from water samples. M.G.C. and F.P.B. were responsible for onboard sampling and generating the environmental dataset. A.T.R.V. was responsible for generating the sequencing dataset. N.M.B., L.N.L., F.V.P., R.G.M.L., D.C.V. and F.M. carried out the bioinformatics and statistical analysis. N.M.B., F.V.P., D.C.V. and F.M. prepared the figures and tables and wrote the first draft of the manuscript. A.G.B. and G.F. provided helpful discussions and technical help. All the authors reviewed the manuscript.

## Acknowledgments

We are grateful to PETROBRAS for the PCR-BS project planning and management. We would like to thank Daniel Moreira for coordinating the PCR-BS project and Dr. Frederico Pereira Brandini for coordinating the “Pelagic realm project”. We would like to thank members of the LECOM team for their scientific support on board. We are deeply grateful for the invaluable guidance and support provided by Rosa Carvalho Gamba; your legacy wisdom continues to inspire us even in her absence. Special thanks to the following PCR-BS project’s researchers and their institutions: UFRJ, UFF, PUC-Rio, UERJ, FIRJAN/SENAI, SALT, INPE, USP, UNIFESP, UFPR, UNESP, IP-SP, FURG and Socioambiental. To the University of São Paulo Foundation (FUSP) for financial administrative management. We also thank the RV Ocean Stalwart and RV Seward Johnson crews and OceanPact for scientific expeditions. We thank Barbara Resende Silva for providing valuable support in designing the conceptual diagram. We acknowledge the anonymous reviewers’ valuable comments, which helped improve this manuscript. We thank Alexandra Lehmkuhl Gerber and Ana Paula C. Guimarães from UGCDFA/LNCC for sequencing the samples.

## Supplementary information

### Additional File 1

Table Supplementary 1. Environmental data of Santos Basin stations from which metagenomes were recovered.

Table Supplementary 2. Metagenomic reads taxonomic annotation at the genus level and its relative abundance in each sample.

Supplementary Table 3. Functional profile of short reads generated through the SEED subsystem and KEGG.

Table Supplementary 4. Post-hoc Dunn test comparing Shannon and Chao1 diversity indices between associations.

Table Supplementary 5. Main features and statistics of 307 Santos Basin metagenome-assembled genomes.

Supplementary Table 6. List of 109 marker genes of the biochemical cycles explored in the Santos Basin Database.

Table Supplementary 7. Metabolic relevant genes present in 153 selected Santos Basin metagenome-assembled genomes.

Table Supplementary 8. Relative abundance of 307 Santos Basin metagenome-assembled genomes at each depth.

### Additional File 2

**Fig. S1.**
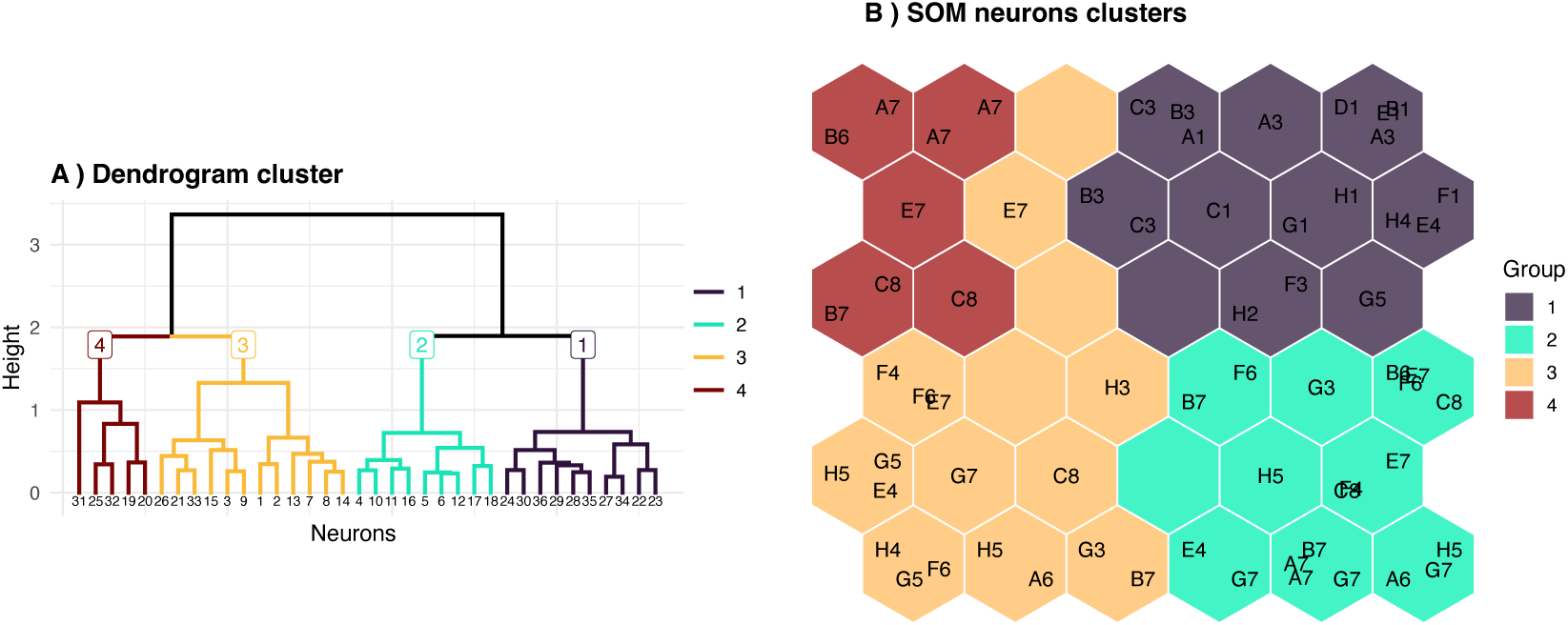
Results from the unsupervised phase of the hybrid model. A: Dendrogram obtained by the hierarchical clustering of the neurons from the self-organizing map (SOM); **B:** SOM with neurons grouped by the respective clusters. The colors represent the associations: association 1 (black), association 2 (cian), association 3 (orange) and association 4 (red).

**Fig. S2.**
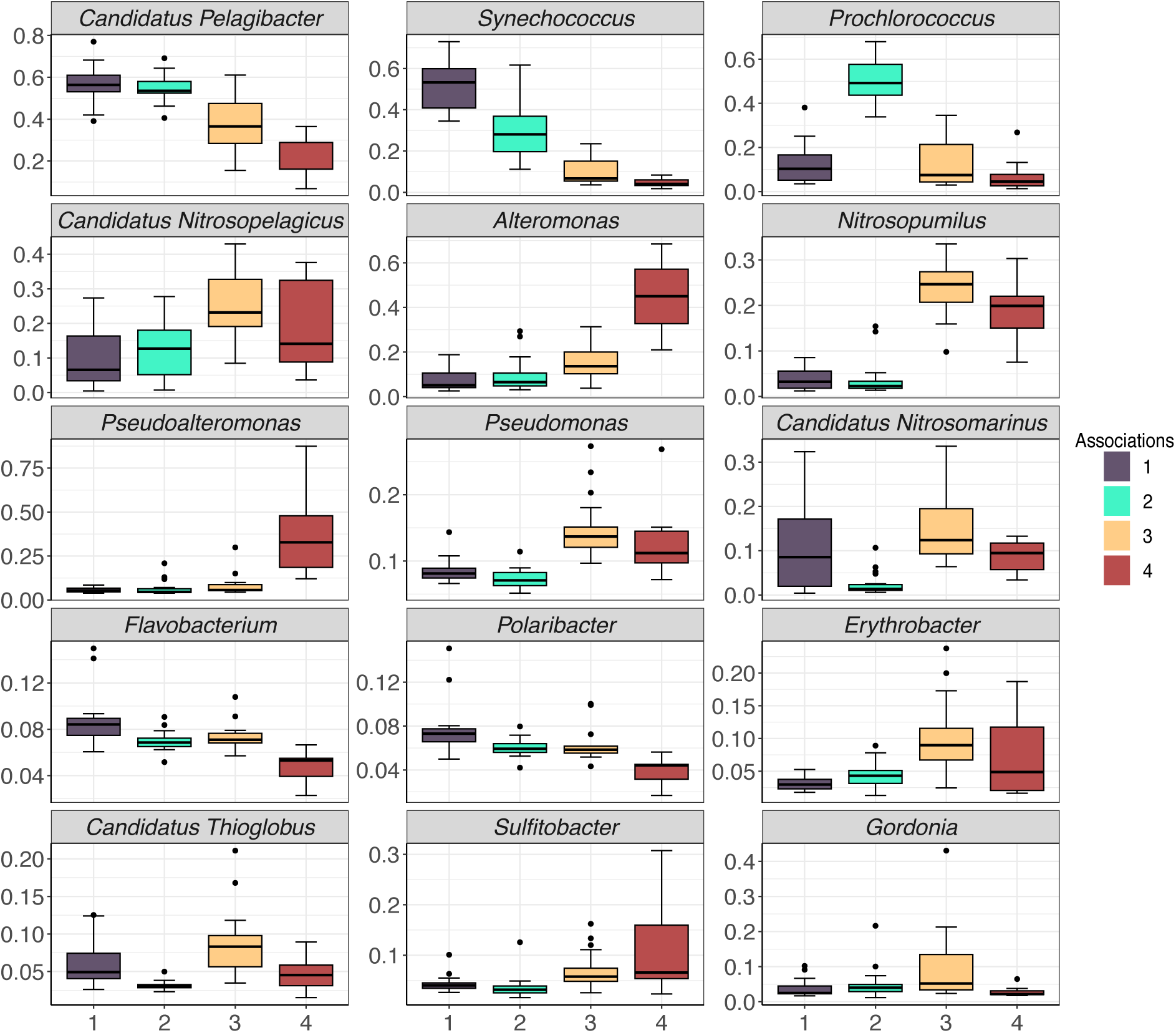
Box plot of the 15 most abundant genera across the four associations. The whiskers represent the minimum and maximum values, the boxes represent the 25th and 75th percentiles, the lines indicate the median values, and the dots represent outliers. The colors represent the associations: association 1 (black), association 2 (cian), association 3 (orange) and association 4 (red).

**Fig. S3.**
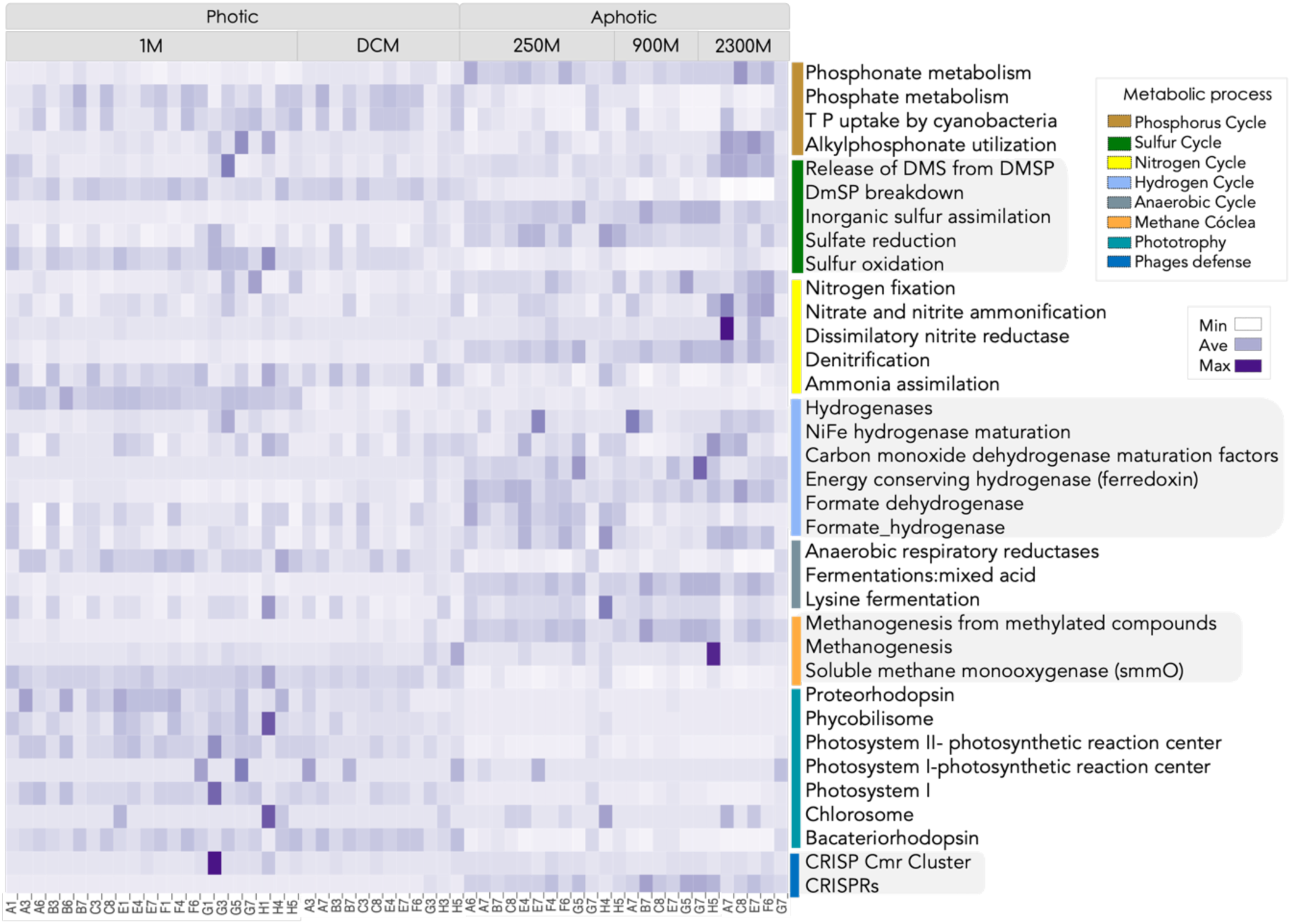
Metabolic profiles obtained from reads, SEED subsystems and pathways, and specific proteins/genes. A blue row Z score scale was applied to statistically assess abundance values differing between samples.

**Fig. S4.**
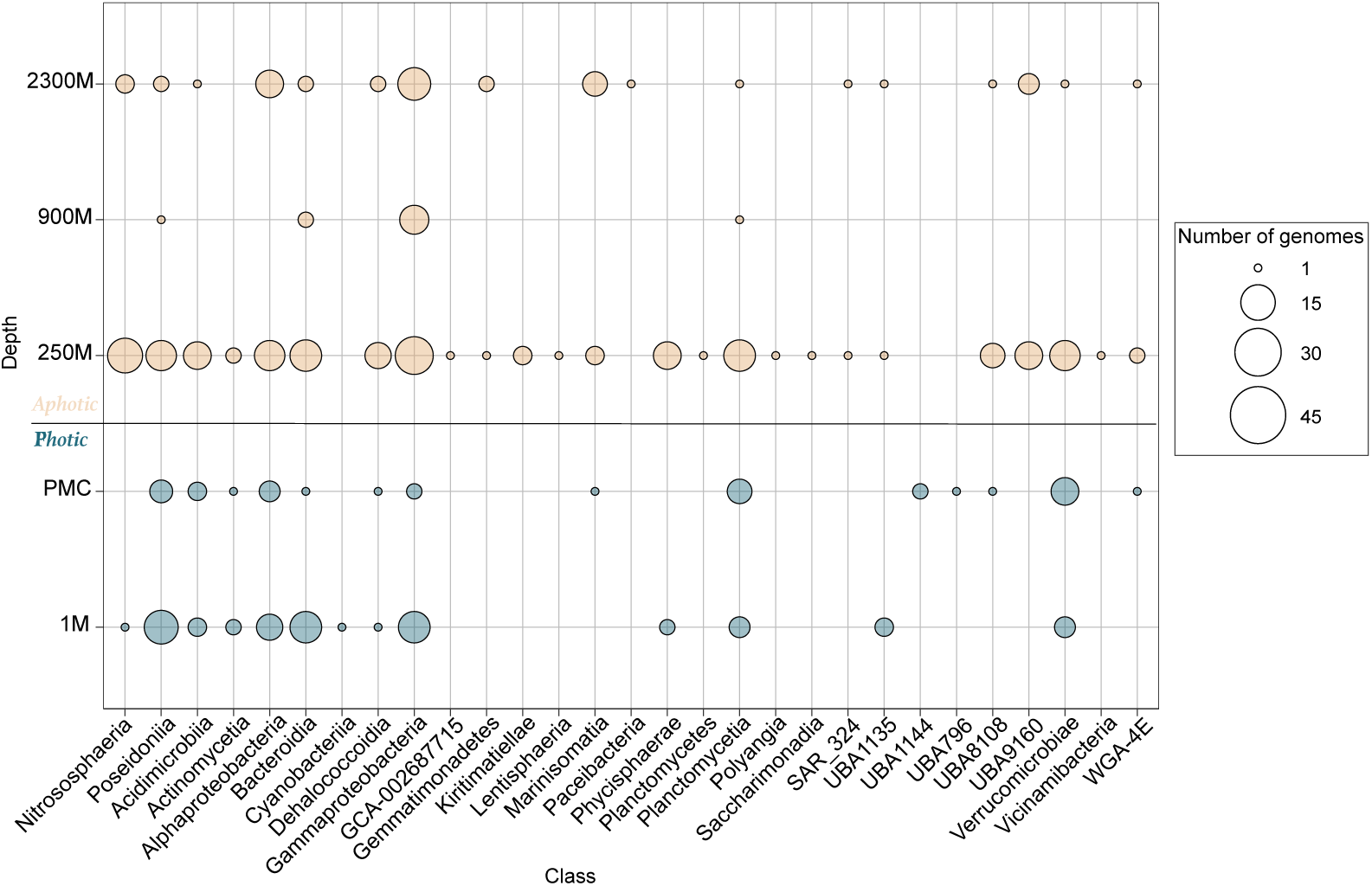
Number of recovered genomes at each depth. MAGs are summarized at the class level, with the size of the circle corresponding to the number of genomes in that class with a given gene, and the color represents the genome from photic (blue) and aphotic (beige) waters. Marker genes that were not detected in any MAG were omitted.

**Fig. S5.**
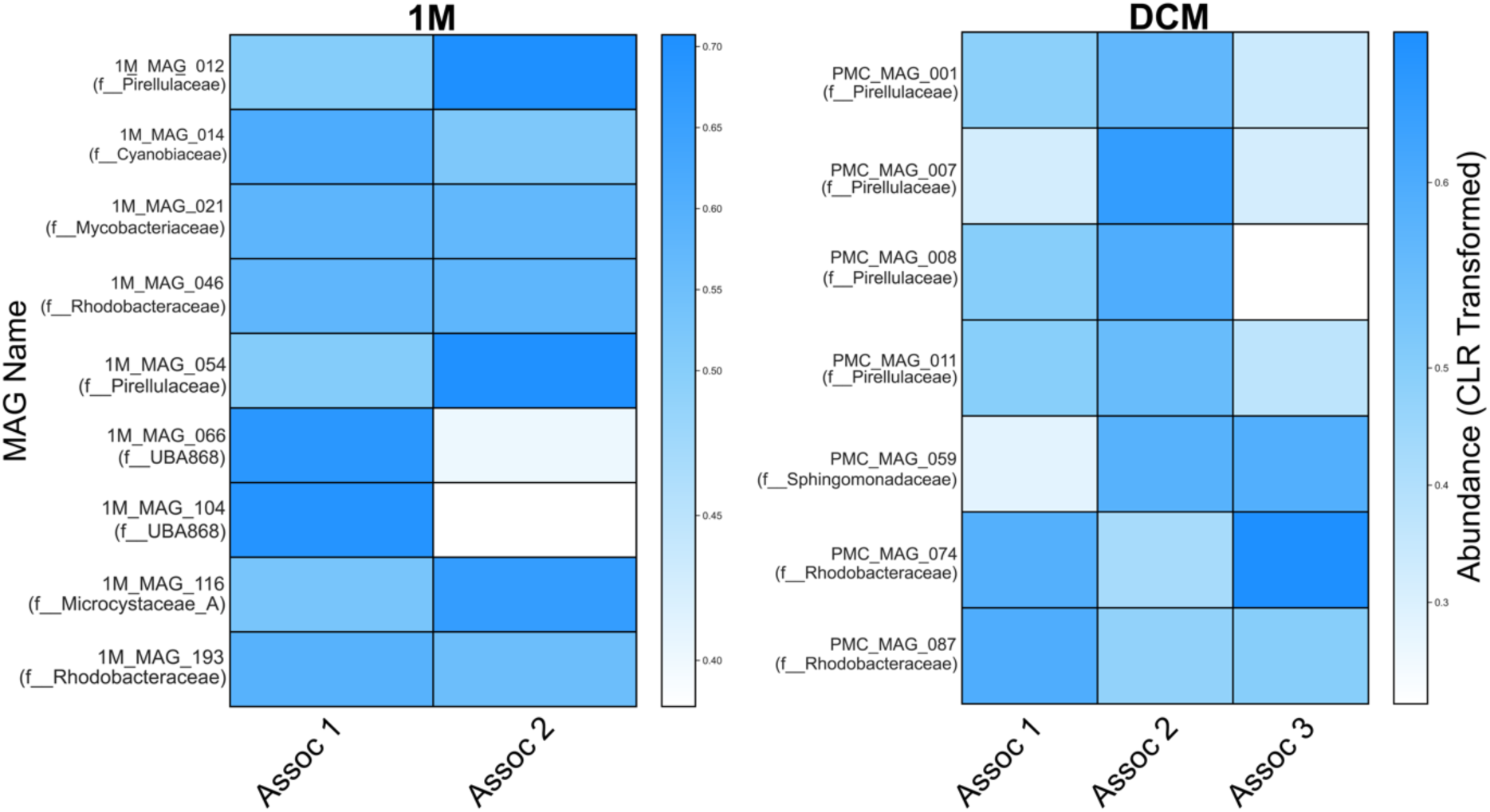
Distribution of metagenome-assembled genomes across SOM associations in the photic environment. The heatmaps display the relative abundances of the genomes at depths of 1 m (left) and DCM (right). Each row represents a MAG, identified by its unique ID and family, whereas columns represent associations (Assoc 1, Assoc 2, and Assoc 3). The color gradient indicates the CLR-transformed genome’s relative abundance values. Each depth was processed independently to highlight depth-specific patterns.

**Fig. S6.**
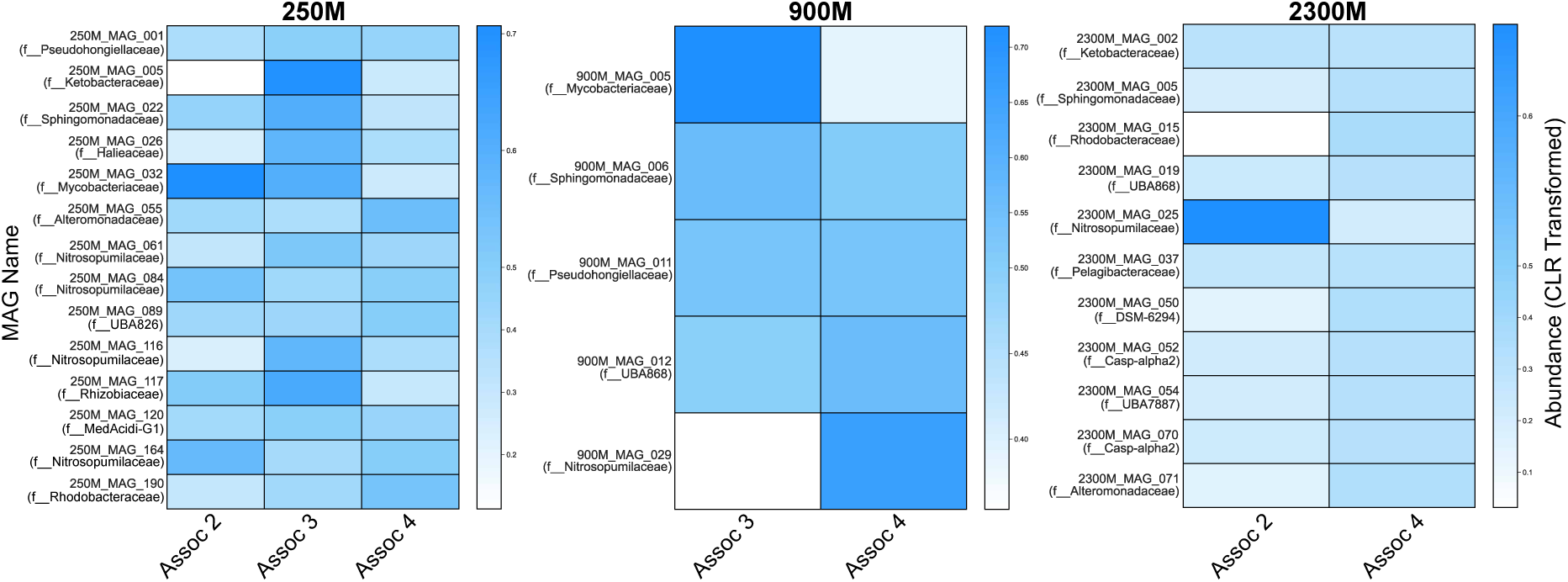
Distribution of metagenome-assembled genomes across SOM associations in the aphotic environment. The heatmaps display the relative abundances of the genomes at depths of 250 m (left), 900 m (center), and 2300 m (right). Each row represents a MAG, labeled with its unique identifier and taxonomic family, whereas columns represent associations (Assoc 2, Assoc 3, and Assoc 4). The color gradient indicates the CLR-transformed genome’s relative abundance values. Each depth was processed independently to highlight depth-specific patterns.

## Notes

### Competing Interest Statement

The authors have declared no competing interest.

